# Mechanistic basis for oxidative stress protection of the human tRNA ligase complex by the oxidoreductase PYROXD1

**DOI:** 10.1101/2023.04.06.535761

**Authors:** Luuk Loeff, Alena Kroupova, Igor Asanović, Franziska Boneberg, Moritz Pfleiderer, Andrè Ferdigg, Fabian Ackle, Javier Martinez, Martin Jinek

## Abstract

RTCB is the catalytic subunit of the metazoan tRNA ligase complex (tRNA-LC) that plays essential roles in tRNA biogenesis and unfolded protein response. The catalytic center of RTCB contains a conserved cysteine that is susceptible to metal ion-induced oxidative inactivation. The flavin-containing oxidoreductase PYROXD1 preserves the activity of mammalian tRNA-LC in a NAD(P)H-dependent manner, but its protective mechanism remains elusive. Here we report a cryo-EM structure of human RTCB in complex with PYROXD1, revealing that PYROXD1 directly interacts with the catalytic center of RTCB through its C-terminal tail. NAD(P)H binding and FAD reduction allosterically control PYROXD1 activity and RTCB recruitment and PYROXD1, while PYROXD1 reoxidation enables timed release of RTCB. PYROXD1 interaction is mutually exclusive with Archease-mediated RTCB guanylylation, and guanylylated RTCB is intrinsically protected from oxidative inactivation. Together, these findings provide a mechanistic framework for the protective function of PYROXD1 that maintains the activity of tRNA-LC under aerobic conditions.

## Introduction

RNA molecules are present in all domains of life and undergo extensive post-transcriptional processing. The precursor transcripts of a subset of tRNA genes contain introns that are removed through a non-canonical spliceosome-independent pathway^1,2^. Non-canonical RNA splicing of precursor-tRNAs (pre-tRNAs) is a two-step enzymatic process that requires endonucleolytic excision of a single intron by the tRNA splicing endonuclease complex^3–5^, after which the two exon halves are ligated by the tRNA ligase complex (tRNA-LC)^6,7^. The core human tRNA-LC is composed of the catalytic subunit RTCB and accessory proteins DDX1, CGI-99, FAM98B, and ASHWIN^6,8,9^. In addition, the complex relies on the co-factor Archease to facilitate multiple turnover catalysis^8^. Besides its essential function in tRNA biogenesis, tRNA-LC is required for XBP1 mRNA splicing in the unfolded protein response^10^, and has been implicated in RNA trafficking^11^ and repair^12^.

RNA ligation by RTCB enzymes proceeds through three distinct nucleotidyl transfer steps. In the first step, catalyzed by the Archease co-factor, a conserved histidine residue in the catalytic center is covalently linked to guanine-5′-monophosphate (GMP) through a nucleophilic attack on the α-phosphate moiety of GTP, releasing inorganic pyrophosphate^13–16^. Subsequently, the 2′,3′-cyclic phosphate of the 5’-exon half is hydrolyzed to yield a 3′-phosphate that executes a second nucleophilic attack on the phosphate group of the RTCB-GMP adduct, resulting in an activated RNA-(3′)pp(5′) G intermediate^13–16^. In the final nucleotidyl transfer step, the 5′-hydroxyl group of the 3′ exon half attacks the activated RNA-(3′)pp(5′)G intermediate of the 5’-exon half, resulting in ligation of the tRNA exon halves and release of GMP^13–16^. Within these catalytic steps, the conserved cysteine in the catalytic center of RTCB coordinates two divalent metal ions that are essential for the ligation reaction^15,16^. Human RTCB contains a strictly conserved catalytic center that comprises a cysteine, an aspartate and four histidine residues that coordinate two divalent metal ions, making RTCB a binuclear metalloenzyme^9^. The reactive cysteine residue in the catalytic center of RTCB renders the enzyme sensitive to oxidative inactivation in the presence of copper(II) ions^17,18^. To preserve the function of RTCB under aerobic conditions, the tRNA-LC relies on the flavin adenine dinucleotide (FAD)-containing oxidoreductase PYROXD1 to protect its catalytic center from oxidation^18^. Depletion of PYROXD1 in cells has been shown to abrogate tRNA ligation, resulting in the accumulation of tRNA exon halves^18^. PYROXD1 oxidizes NAD(P)H to NAD(P)^+^ in a tightly controlled manner, sustaining the ligase activity of the tRNA-LC under oxidizing conditions in vitro^18^. FAD-bound PYROXD1 directly interacts with RTCB in an NAD(P)H-dependent manner^18^; however, the mechanistic basis for the protective function of PYROXD1 has remained elusive.

Here, we present a cryogenic electron microscopy (cryo-EM) structure of human RTCB in complex with PYROXD1 and reveal the molecular basis for PYROXD1-mediated protection of RTCB. Our results demonstrate that the C-terminal tail of PYROXD1 directly interacts with the catalytic center of RTCB and identify a NAD(P)H-dependent conformational switch in PYROXD1 that allosterically modulates the interaction with RTCB. We further show that guanylylation of the catalytic center of RTCB by Archease inhibits the recruitment of PYROXD1 and intrinsically protects RTCB from oxidative inactivation. Our results collectively reveal how PYROXD1 protects the catalytic subunit of the tRNA-LC from inactivation by reactive oxygen species.

## Results

### Molecular architecture of the RTCB-PYROXD1 complex

To elucidate the mechanism underlying the protective function of the flavoprotein PYROXD1 in preventing oxidative inactivation of RTCB, we first used affinity co-precipitation experiments to establish the conditions under which RTCB and PYROXD1 stably interact. In agreement with prior studies^18^, recombinant PYROXD1 was co-precipitated by immobilized RTCB in an NADH-dependent manner, requiring the presence of 0.5 mM NADH for a stable, stoichiometric interaction (**Figure S1A**). PYROXD1 co-precipitation was observed in the presence of a variety of divalent metal cations (Ca^2+^, Co^2+^, Cu^2+^, Mg^2+^, Mn^2+^, Ni^2+^ & Zn^2+^), while chelation of divalent cations with ETDA abolished the interaction between PYROXD1 and RTCB (**Figure S1B**), confirming that the interaction is divalent cation-dependent^18^. As monitored by absorption spectroscopy, PYROXD1 catalyzed multiple-turnover oxidation of NADH to NAD^+^ (**Figure S1C**); the observed kinetics are consistent with previous studies showing that PYROXD1 catalyzes the rapid initial formation of a NAD^+^:FADH^-^ charge-transfer complex (CTC), followed by slow reoxidation of the CTC by molecular O^_2_ 18^.

To visualize the interaction between RTCB and PYROXD1, we reconstituted the RTCB-PYROXD1 complex in the presence of NADH and Mg^2+^ ions, and analyzed it using single-particle cryo-EM. To overcome the orientation bias of the particles, two datasets were acquired using samples prepared in the presence and absence of octyl-beta-glucoside (**Figure S2A**), and combined to yield a molecular reconstruction of the RTCB-PYROXD1 complex at a resolution of 3.3 Å (**Figure 1A-B** and **S2B-C**). The resulting model is comprised of a single molecule of RTCB and PYROXD1, indicating that RTCB and PYROXD1 interact with a 1:1 stoichiometry (**Figure 1A-B**). The interaction involves the C-terminal domain (CTD) of PYROXD1, whose function has hitherto remained unclear, and the active site cleft of RTCB, spanning an extensive interface with a buried surface area of 1715 Å^2^.

**Figure 1:**
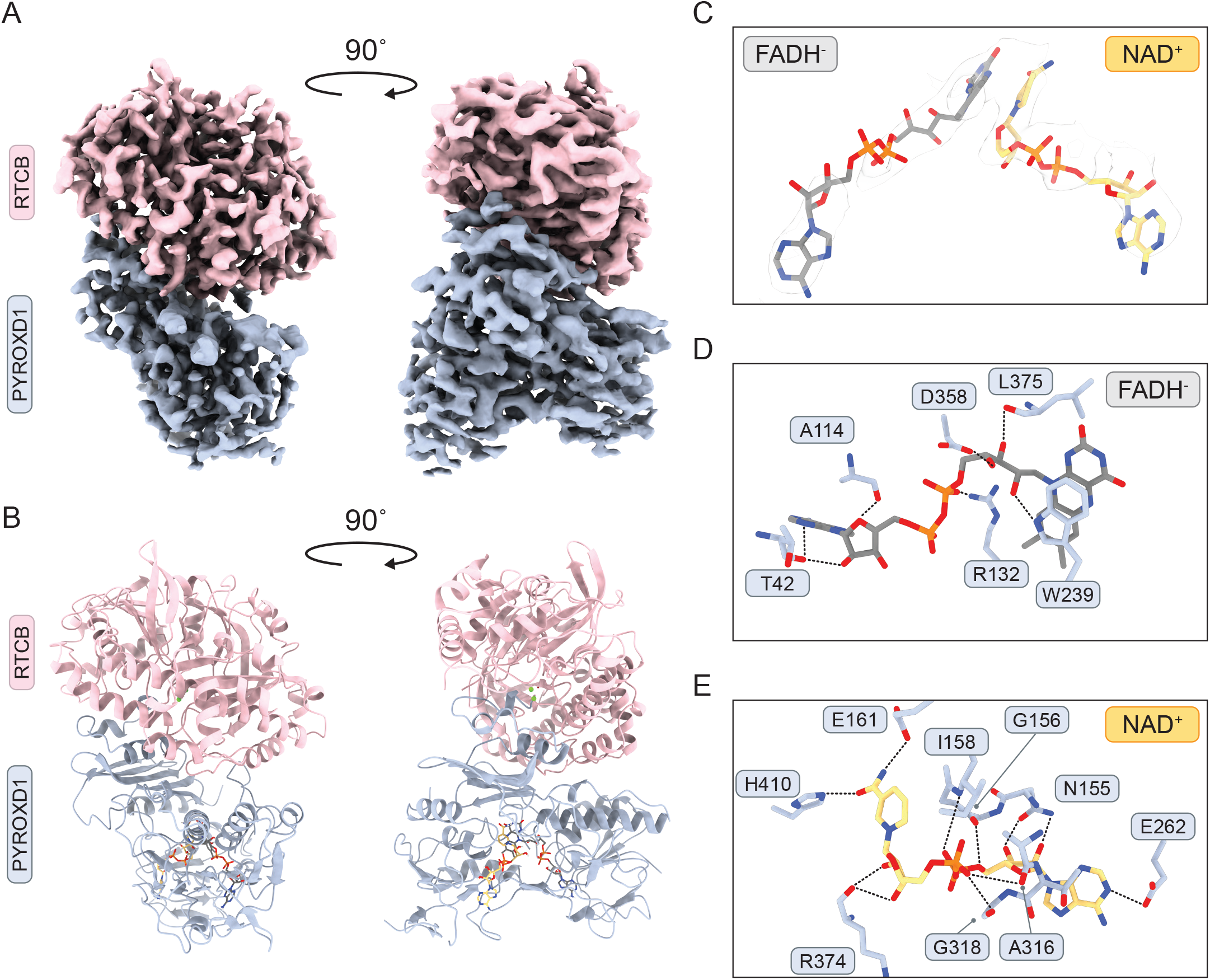
Cryo-EM structure of the human RTCB-PYROXD1 complex. **(A)** Cryo-EM density map of the RTCB-PYROXD1 complex. **(B)** Structural model of the RTCB-PYROXD1 complex. **(C)** FADH-(dark grey) and NAD+ (orange) ligands in the RTCB-PYROXD1 complex. Map (light grey) contoured at s=10. **(D)** Interaction map of the FADH-ligand in the RTCB-PYROXD1 complex. **(E)** Interaction map of the NAD+ ligand in the RTCB-PYROXD1 complex.

The molecular reconstruction shows clear densities for the nicotinamide and flavin dinucleotides within the catalytic center of PYROXD1 (**Figure 1C**). Based on the kinetics of NADH oxidation and the NAD(P)H dependence of the RTCB-PYROXD1 interaction, we infer that the PYROXD1 adopts the NAD^+^:FADH^-^ CTC state when bound to RTCB. As in other flavoprotein oxidoreductases^19–24^, the ligands are positioned by PYROXD1 such that the flavin ring of the FADH^-^ molecule stacks with the nicotinamide ring of NAD^+^, forming a pi-pi interaction (**Figure 1C**). PYROXD1 coordinates FADH^-^ with residues Thr42^PYROXD1^, Ala114^PYROXD1^, Arg132^PYROXD1^, Trp239^PYROXD1^, Asp358^PYROXD1^ and Leu375^PYROXD1^ (**Figure 1D** and **Figure S3A**). In turn, NAD^+^ interacts with residues Asn155^PYROXD1^, Gly156^PYROXD1^, Ile158^PYROXD1^, Glu161^PYROXD1^, Ala316^PYROXD1^, Gly318^PYROXD1^, Arg374^PYROXD1^ and His410^PYROXD1^ (**Figure 1E** and **Figure S3B**). In sum, these structural and biochemical observations suggest that the interaction of PYROXD1 with RTCB is dependent on NAD(P)H binding in the PYROXD1 active site center and the resulting formation of the CTC.

#### C-terminal tail of PYROXD1 protects the RT catalytic center

The structure of the RTCB-PYROXD1 complex reveals that the PYROXD1 CTD engages with the catalytic center cleft of RTCB, thereby blocking the substrate binding sites and rendering the catalytic center inaccessible (**Figure 2A**). The interaction is centered on the alpha-helical C-terminal tail of PYROXD1 (residues Glu496–Asp500), which insets into the catalytic center of RTCB and forms additional hydrophobic interactions with RTCB via Ile394^PYROXD1^, Ile395^PYROXD1^, Tyr498^PYROXD1^ and Phe499^PYROXD1^ (**Figure 2B**). The C-terminal helix of PYROXD1 is anchored in the RTCB catalytic center by hydrogen bonding interactions of the C-terminal carboxyl group with the backbone amide groups of Phe118^RTCB^ and Asp119^RTCB^. Furthermore, the side chain of Asp500^PYROXD1^ coordinates one of the two Mg^2+^ ions (metal position B) in the RTCB catalytic center, in agreement with the observation that the PYROXD1-RTCB interaction is dependent on the presence of divalent cations (**Figure S1B**).

**Figure 2:**
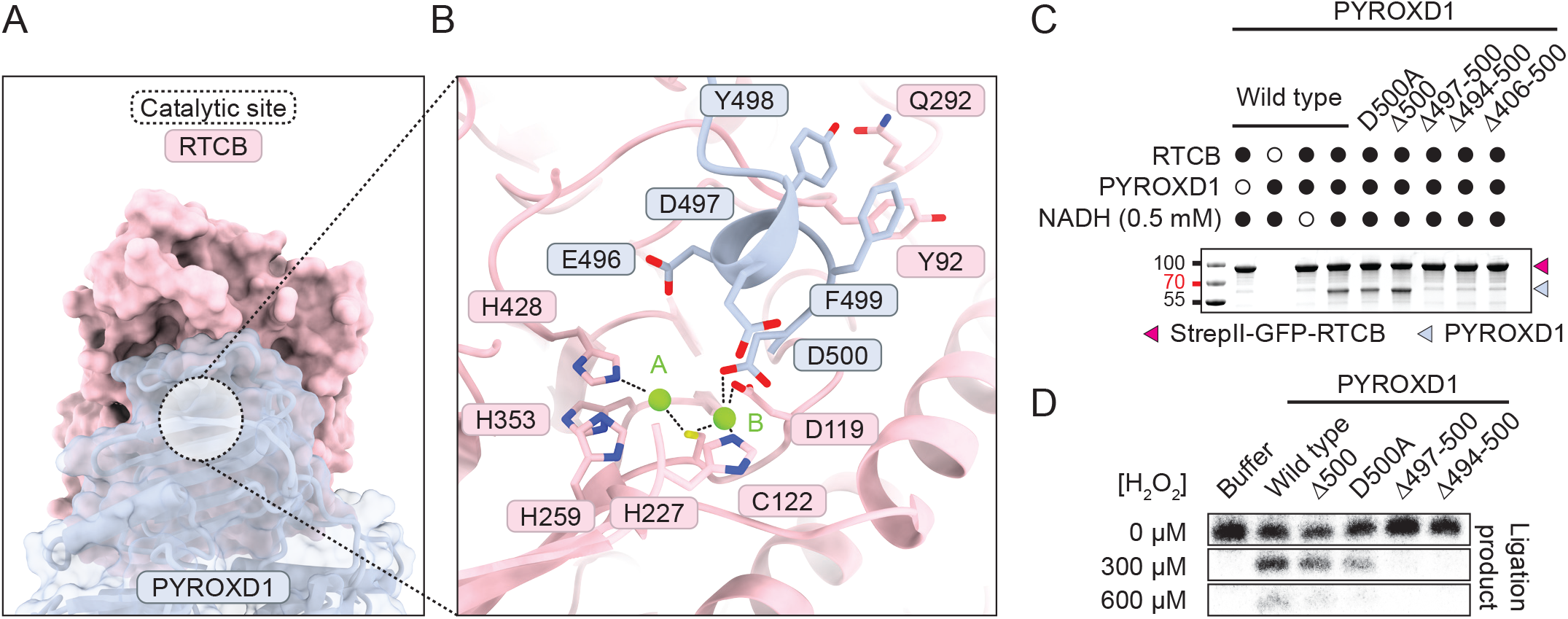
The C-terminal domain of PYROXD1 blocks the catalytic center of RTCB. **(A)** Surface view of the RTCB-PYROXD1 interaction. The catalytic site of RTCB is highlighted by the dashed circle. **(B)** Detailed view of the interaction of the CTD of PYROXD1 with the catalytic site of RTCB. **(C)** In vitro pull-down experiment with recombinant PYROXD1 mutants and StrepII-GFP-RTCB. Strep-Tactin beads were washed to remove unbound PYROXD1, and bound proteins were analyzed by SDS-PAGE and Coomassie blue staining. **(D)** In vitro ligation assay in presence and absence of oxidative stress.

To validate the structural determinants of the interaction, we generated PYROXD1 protein constructs containing either an alanine substitution or deletion of the C-terminal residue (Asp500^PYROXD1^) or truncations (Δ497-500, Δ494-500, Δ406-500) of the CTD, and performed in vitro affinity co-precipitation experiments using immobilized RTCB. While PYROXD1 proteins containing the D500A^PYROXD1^ mutation or lacking the C-terminal residue retained their ability to interact with RTCB, PYROXD1 constructs containing more extensive C-terminal truncations of the CTD were not co-precipitated by RTCB (**Figure 2C**), confirming that the C-terminal alpha-helix is required for the interaction. Together, these results reveal that the C-terminal domain of PYROXD1, and specifically its C-terminal helix, are critical for the divalent cation-dependent interaction of PYROXD1 with RTCB.

Next, we tested the importance of the RTCB interaction interface in PYROXD1 for the protective function of PYROXD1 against oxidative inactivation of RTCB. To this end, we performed an RTCB RNA circularization assay upon incubating RTCB in the presence of wild-type (WT) and mutant PYROXD1 proteins and increasing concentrations of hydrogen peroxide^18^. In the absence of hydrogen peroxide, RTCB efficiently catalyzed RNA circularization in all reactions. By contrast, RTCB only remained catalytically active upon hydrogen peroxide treatment in the presence of PYROXD1 protein variants capable of interacting with RTCB; RTCB activity was not sustained by C-terminally truncated PYROXD1 proteins that did not interact with RTCB (**Figure 2D**). Notably, RTCB activity was reduced upon peroxide treatment in the presence of PYROXD1 mutants Δ500 and D500A^PYROXD1^, as compared to WT PYROXD1 (**Figure 2D**), suggesting that these PYROXD1 mutants bound to RTCB with a reduced affinity, possibly due to impaired metal ion coordination. Taken together, these results confirm that the interaction of the C-terminal tail of PYROXD1 with the catalytic center of RTCB is essential for the protection of RTCB from oxidative inactivation.

#### Allosteric mechanism of the RTCB-PYROXD1 interaction

Superposition of the RTCB-PYROXD1 complex structure with the crystal structure of NAD(P)H-free PYROXD1^18^ reveals structural rearrangements in PYROXD1 induced by NADH binding and subsequent CTC formation and RTCB recruitment. A conserved loop (residues Pro45^PYROXD1^-Arg76^PYROXD1^) within the FAD binding domain of PYROXD1, hereafter referred to as Loop 1, is positioned away from FAD in the PYROXD1 crystal structure^18^. In the RTCB-PYROXD1 complex, Loop1 is only partially ordered and shifted towards the active center cavity that houses the NAD^+^:FADH^-^ CTC (**Figure 3A**). In addition, this rearrangement positions the loop such that it avoids a steric clash with RTCB and instead interfaces with it via a strictly conserved sequence motif (Motif 1) (**Figure S3C**). However, its functional significance for RTCB binding could not be tested as point mutations in Motif 1 or deletion of entire Loop1 render the mutant proteins insoluble, presumably by compromising the ability of PYROXD1 to bind FAD.

**Figure 3:**
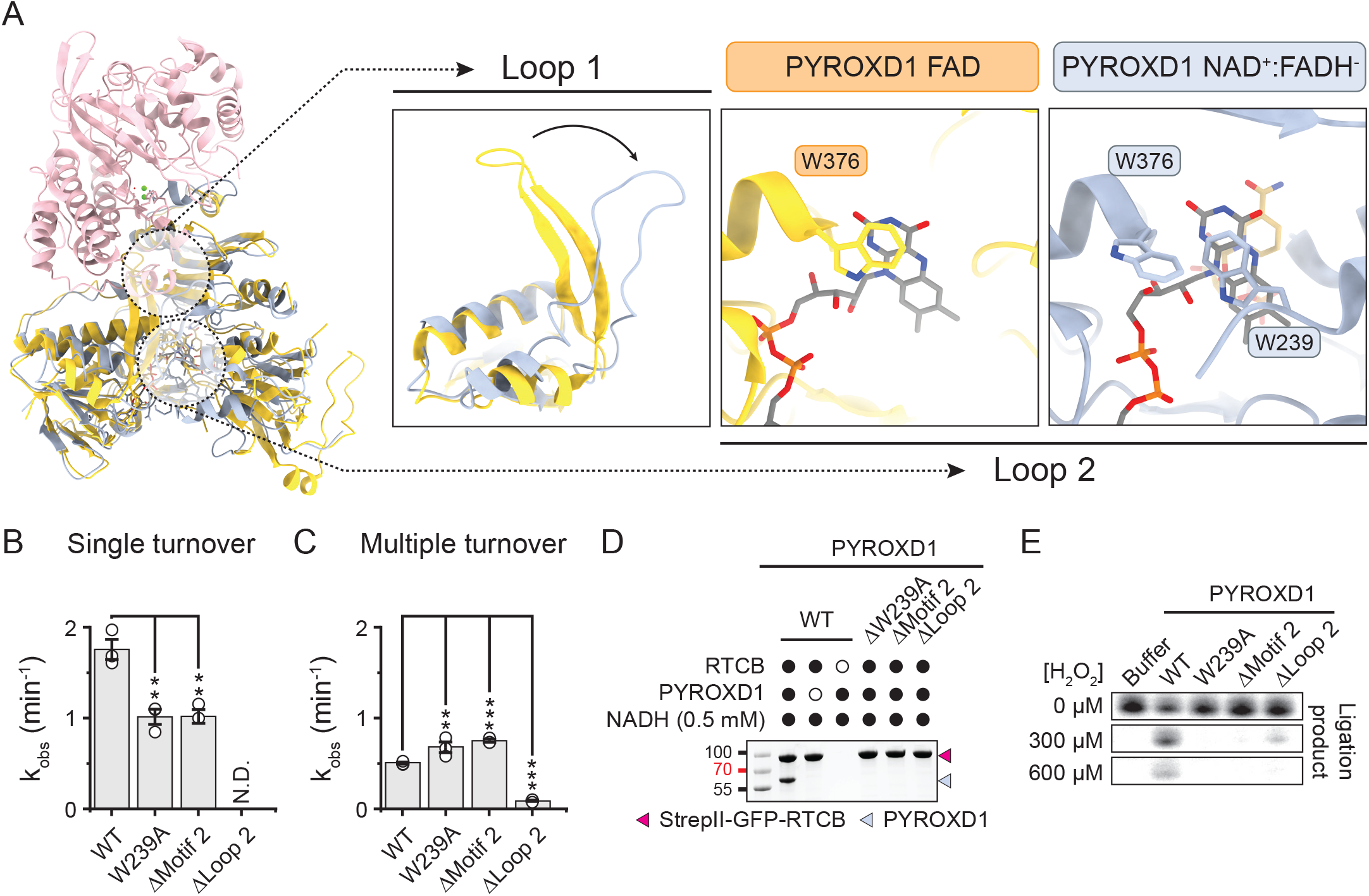
Allosteric regulation of PYROXD1 mediates CTC formation and turnover. **(A)** Structural super position of PYROXD1 bound to FAD (orange, PDB: 6ZK7)18 and PYROXD1 (light blue) in complex with RTCB (pink). **(B)** Kinetic rates for NADH turnover by PYROXD1 wild-type and Loop 2 mutants under pre-steady state conditions. Data points represent the mean ± SEM of three independent replicates. Significance was determined by using an unpaired two-tailed t-test. P-values: ** denotes p ≤ 0.01. **(C)** Kinetic rates for NADH turnover by PYROXD1 wild-type and Loop 2 mutants under steady state conditions. Data points represent the mean ± SEM of three independent replicates. Significance was determined by using an unpaired two-tailed t-test. P-values: ** denotes p ≤ 0.01, *** denotes p ≤ 0.001. **(D)** In vitro pull-down experiment with recombinant PYROXD1 mutants and StrepII-GFP-RTCB. Strep-Tactin beads were washed to remove unbound PYROXD1, and bound proteins were analyzed by SDS-PAGE and Coomassie blue staining. **(E)** In vitro ligation assay in presence and absence of oxidative stress.

Additionally, a long disordered loop (residues Ala190^PYROXD1^-Glu250^PYROXD1^) that connects the NAD(P)H binding domain and the CTD of PYROXD1, hereafter referred to as Loop 2, becomes partially ordered in the RTCB-PYROXD1 complex (**Figure 3A**). The loop is channeled towards the NAD^+^:FADH^-^ ligands in the complex, positioning a conserved motif (Motif 2) such that the side chain of Trp239^PYROXD1^ can interact with the flavin ring of FADH^-^ (**Figure 3A** and **Figure S3D**). As a consequence, Trp376^PYROXD1^, which caps the flavin ring of FAD in the NAD(P) H-free PYROXD1^18^ and limits its access to electron acceptors, is displaced away in the RTCB-PYROXD1 complex (**Figure 3A**). These observations suggest that Loop 2 modulates the NAD(P)H oxidase activity of PYROXD1 by regulating access to electron acceptors.

To validate the importance of Loop 2, we generated mutant PYROXD1 proteins containing a point mutation in Motif 2 (W239A^PYROXD1^), deletion of Motif 2 (ΔMotif 2) or a deletion of the entire Loop 2 (ΔLoop 2) and measured their catalytic rates of NADH oxidation. Under single-turnover conditions, which reflected CTC formation, mutations of Motif 2 (W239A^PYROXD1^ and ΔMotif 2) resulted in ∼2-fold slower rates of NADH oxidation (**Figure 3B** and **S3E**). By contrast, under steady state conditions, which reflected turnover of the CTC, mutations of Motif 2 (W239A^PYROXD1^and ΔMotif 2) resulted in ∼1.5-fold faster rate of NADH oxidation (**Figure 3C** and **S3F**). In contrast, the ΔLoop 2 PYROXD1 mutant lost its ability to catalyze NADH oxidation (**Figure 3B-C**). Co-precipitation experiments revealed that the mutations of Motif 2 or deletion of Loop 2 abolished interactions of PYROXD1 with RTCB (**Figure 3D**). In agreement with these results, both the Motif 2 and Loop 2 deletion PYROXD1 mutants failed to protect RTCB from oxidative inactivation in vitro (**Figure 3E**). In sum, these data indicate that Loop 2 allosterically modulates the oxidoreductase and RTCB binding activities of PYROXD1, thereby underpinning its protective function against oxidative inactivation of RTCB.

#### PYROXD1 protects RTCB prior to Archease mediated guanylylation

PYROXD1 sterically blocks the RTCB catalytic center cleft, as supported by superposition of the RTCB-PYROXD1 complex structure with the structure of *Pyrococcus horikoshii* RtcB bound to a 5’-hydroxyl DNA oligonucleotide^25^ (**Figure S4A**), suggesting that the PYROXD1-RTCB interaction precludes substrate binding and catalytic activity of RTCB. In turn, close inspection of a superposition of the complex with the crystal structure of human RTCB bound to GMP indicates steric clashes of PYROXD1 with a loop comprising residues Asp444-Glu474^RTCB^, which is structurally disordered in the PYROXD1-RTCB complex but adopts a defined conformation in the RTCB-GMP structure due to interactions with bound GMP (**Figure S4B-E**). Additionally, binding of PYROXD1 to guanylylated RTCB would result in a steric clash between the side chain of Asp497^PYROXD1^ and the alpha-phosphate group of GMP (**Figure S4B-E**). Together, these structural observations thus suggest that PYROXD1 selectively interacts with the apoenzyme form of RTCB, that is, prior to guanylylation of the catalytic center histidine His428^RTCB^.

To test whether PYROXD1 interacts with the guanylylated or un-guanylylated forms of RTCB, we first preincubated RTCB in the presence of GTP, Mn^2+^ ions and human Archease, generating a mixture of guanylylated and un-guanylylated RTCB. We subsequently performed a co-precipitation assay using immobilized PYROXD1, and analyzed the unbound and co-precipitated RTCB fractions by mass spectrometry. Guanylylated RTCB was substantially depleted from the co-precipitated (eluted) fraction, as compared to the unbound fraction and input fractions, suggesting that PYROXD1 preferentially associates with un-guanylylated, apo-RTCB (**Figure 4A** and **S4F-G**). As this implies that guanylylated RTCB cannot be protected by PYROXD1, we followed up on this observation and tested whether guanylylated RTCB is susceptible to oxidative inactivation. In this experiment, apo-RTCB or pre-guanylylated RTCB were treated with hydrogen peroxide in the presence or absence of PYROXD1, and subsequently assayed for RNA ligase activity (**Figure 4B**). In the absence of pre-guanylylation, RTCB was inactivated by hydrogen peroxide, which could be mitigated by addition of PYROXD1 during peroxide treatment. In contrast, pre-guanylylated RTCB remained partially catalytically active upon hydrogen peroxide treatment in the absence of PYROXD1, suggesting it is considerably less susceptible to peroxide-induced oxidative inactivation (**Figure 4B** and **S4H**). Together, these results suggest that PYROXD1 protects apo-RTCB, while guanylylated RTCB does not interact with PYRODX1 and is intrinsically protected oxidative inactivation.

**Figure 4:**
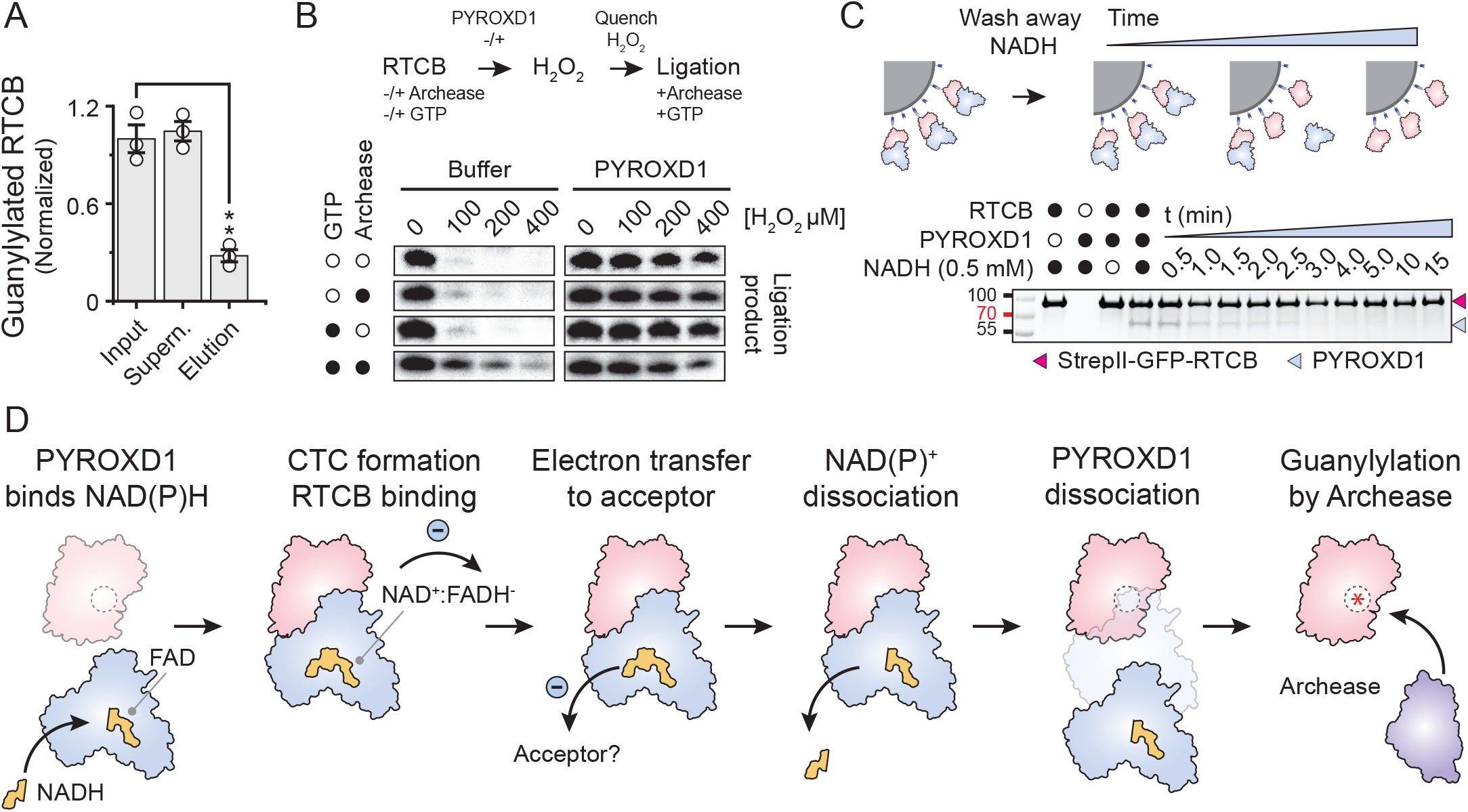
Mechanism for oxidative stress protection of tRNA ligase by the oxidoreductase PYROXD1. **(A)** Normalized guanylylation levels of RTCB determined by LC-MS in the input, supernatant and elution samples of a pulldown with GST-PYROXD1. Data points represent the mean ± SEM of three independent replicates. Significance was determined by using an unpaired two-tailed t-test. P-values: ** denotes p ≤ 0.01. **(B)** In vitro ligation assay to determine the effects of guanylylation on the oxidative inactivation of RTCB. The ligation assay was performed in a three-step procedure, where RTCB was incubated with various combinations of GTP and Archease (see labels on the left), followed by incubation with various concentrations of H_2_O_2_ in the presence or absence of PYROXD1. In the last steps all samples were supplemented with GTP and Archease to allow for multiple turn-over ligation of the RNA substrate. Panels are a composite of separate experiments, see Figure S4H for individual gel panels. **(C)** In vitro pull-down experiment with recombinant PYROXD1 and StrepII-GFP-RTCB. Strep-Tactin beads were washed to remove unbound PYROXD1 and NADH from solution. Next, at the indicated times, bound proteins were analyzed by SDS-PAGE and Coomassie blue staining. **(D)** Model for the protection of RTCB by PYROXD1 against oxidative stress. First, PYROXD1 binds to FAD and NAD(P)H and forms a CT complex by transferring an electron from NAD(P)H to FAD. The CT complex is recruited to the catalytic center of RTCB, occluding the catalytic center from the solvent. PYROXD1 transfers an electron to an acceptor, such as reactive oxygen, which results in the release of NAD+. When NAD+ is released PYROXD1 dissociates from RTCB, allowing Archease to access and guanylylate the catalytic center for multiple turn-over ligation of pre-tRNAs.

#### PYROXD1 reoxidation controls RTCB release

The initial step of RTCB-catalyzed RNA ligation involves Archease-mediated guanylylation of the RTCB catalytic center. As PYROXD1 occludes the catalytic center of RTCB, dissociation of the RTCB-PYROXD1 complex is thus required prior to RTCB guanylylation and subsequent RNA ligation. In a co-precipitation experiment, Archease was not able to displace PYRODX1 from RTCB even when supplied in a large molar excess together with GTP (**Figure S4**I). This result is consistent with previous observations that Archease does not interact stably with RTCB or tRNA-LC, indicative of a transient interaction^9,27^. We hypothesized that instead of direct competition with Archease, the dissociation of the RTCB-PYROXD1 complex is triggered by reoxidation of the NAD^+^:FADH^-^ CTC in PYROXD1 by an electron acceptor (such as molecular oxygen), resulting in FAD regeneration and NAD^+^ release. To test this, we analyzed the dissociation kinetics of the RTCB-PYROXD1 complex in vitro under aerobic conditions. The complex was reconstituted in the presence of 0.5 mM NADH and immobilized on an affinity (StrepTactin) matrix via RTCB. Excess NADH was washed away and loss of PYROXD1 co-precipitation was monitored over time (**Figure 4C**). The estimated half-life of the RTCB-PYROXD1 upon NADH removal was ∼2 min, comparable to the rate of PYROXD1 CTC reoxidation (**Figure 3C** and **4C**). This suggests that PYROXD1 rapidly dissociates from RTCB upon reoxidation by an electron acceptor. Taken together, these data suggest a mechanism for the timed release of RTCB, enabling its catalytic activation by Archease.

## Discussion

The RNA ligase activity of tRNA-LC is essential for cellular RNA metabolism. The enzymatic mechanism of the catalytic subunit RTCB has anaerobic origins^15,16^ and is dependent on a highly reactive cysteine residue, which makes the enzyme sensitive to inactivation under oxidative stress^17,18^. In metazoa, RTCB has co-evolved with the oxidoreductase PYROXD1 that protects RTCB from oxidative inactivation, while in plants and fungi RTCB has been functionally replaced by an evolutionarily unrelated RNA ligase with a distinct catalytic mechanism^18,26^. In this work, we employed single-particle cryo-EM combined with biochemical experiments to provide a mechanistic basis for the protective function of PYROXD1. Our results further underscore the notion of co-evolution of PYROXD1 with RTCB as a mechanism to enable the tRNA-LC to function under aerobic conditions. Although RTCB was previously suggested to bind NAD(P)H directly^18^, our structural study reveals that NAD(P)H is instead required for allosteric conformational activation of PYROXD1 to enable RTCB binding via an extensive molecular interface involving the C-terminal tail of PYROXD1 as well as divalent cations in the RTCB catalytic center. Based on these results, we propose a model in which PYROXD1 protects RTCB from reactive oxygen species by physically occluding its catalytic center (**Figure 4D**) and inhibiting metal ion exchange, thereby preventing oxidation of the cysteine in the catalytic center by excluding Cu^2+^ ions and reactive oxygen species.

PYROXD1 undergoes a conformational change upon binding of NAD(P)H, which facilitates the electron transfer from NAD(P)H to the FAD co-factor and results in the formation of a CTC (**Figure 4D**). This conformational rearrangement positions two loops of PYROXD1 into the vicinity of the NAD^+^:FADH^-^ ligands within the protein allowing for interaction with RTCB and controlled turnover of the CTC. Consistent with this model, our biochemical data show that loop mutations that compromise the formation of the CTC or increase its turnover impair RTCB binding and the protective function of PYROXD1.

The catalytic activity of RTCB is dependent on the co-factor Archease, which catalyzes the initial guanylylation of a conserved histidine residue in the RTCB catalytic center. A recently determined structure of the RTCB-Archease complex reveals the mechanism of Archease-assisted guanylylation^27^ and indicates that Archease and PYROXD1 interact with RTCB in a mutually exclusive manner. This is consistent with our observations that PYROXD1 selectively interacts with RTCB in the apo-enzyme form prior to guanylylation, and that guanylylated RTCB is intrinsically protected from oxidative inactivation. Overall, our results suggest that the NAD(P)H oxidase activity of PYROXD1 serves as a molecular timer for controlled release of RTCB for Archease-mediated activation, thereby enabling tRNA-LC function under oxidative stress conditions^2,18^. As the coupling of RTCB protection to the oxidoreductase activity of PYROXD1 also likely serves a regulatory function, this study establishes a paradigm for redox-dependent control of RNA processing pathways.

## Data and code availability

Atomic coordinates and cryo-EM maps for the human RTCB-PYROXD1 complex (PDB: xxx), EMDB: EMD-xxx), have been deposited in the PDB and EMDB databases and are publicly available as of the date of publication. Accession codes are listed in the key resources table. Any additional information required to reanalyze the data reported in this paper is available from the lead contact upon request.

## Acknowledgements

We would like to thank Marta Sawicka, Simona Sorrentino and the UZH Center for Microscopy and Image Analysis for technical support with cryo-EM data acquisition. We thank Jirka Peschek and Janina Gerber for fruitful discussions and sharing unpublished data. We thank Scott Gradia (UC Berkeley MacroLab) for providing LIC cloning vectors. We thank Serge Chesnov from the functional genomics center in Zurich for assistance with processing and data acquisition of LC-MS samples. LL was funded by a Horizon 2020 Marie Skłodowska-Curie Individual Fellowship (project no. 845268, MSOPGDM). AK was funded by the Boehringer Ingelheim Fonds PhD Fellowship and the Forschungskredit program of the University of Zurich (grant no. FK-18-033). MJ is International Research Scholar of the Howard Hughes Medical Institute, Vallee Scholar of the Vallee Foundation and member of the Swiss National Competence Center for Research “RNA & Disease”.

## Author Contributions

AK, IA, JM, LL, MJ conceived study. AK, FA, FB, IA, LL, MP purified proteins. LL collected and processed cryo-EM data. AK, FA, FB, IA, LL, MP performed biochemical experiments. LL, and MJ wrote manuscript with input from all authors.

## Methods

### Plasmid DNA constructs and site-specific mutants

DNA fragments encoding human Archease (Uniprot Q8IWT0), human PYROXD1 (Uniprot Q8WU10) and human RTCB (UniProt Q9Y3I0) were obtained as described previously^9,18^. In brief, human PYROXD1 was amplified from HeLa cDNA by polymerase chain reaction (PCR) using primers BamHI_PYROXD1_F (5’-CGC GGA TCC ATG GAG GCA GCG CGC CCT CC-3’) and XhoI_PYROXD1_R (5’-TAG CCG CTC GAG TTA GTC AAA ATA ATC TTC TA-3’). The amplified sequence was ligated into the pGEX-6P-1 vector (GE Healthcare) upon cleavage with the restriction enzymes BamHI and XhoI. The resulting construct carries an N-terminal Glutathione tag followed by a 3C Precision protease cleavage site. Mutants of PYROXD1 were generated by Gibson assembly^30^, using gBlocks synthesized by IDT. The human Archease gene spanning residues 27-168 was inserted into the 2M-T (Addgene: 29708) plasmid using ligation-independent cloning (LIC), resulting in a construct carrying an N-terminal hexahistidine tag followed by a maltose binding protein (MBP) and a tobacco etch virus (TEV) protease cleavage site. The human RTCB gene was inserted into the 438B (Addgene: 30115) plasmid and the 438-Rgfp (Addgene: 55221) plasmid, using LIC. The resulting constructs carries an N-terminal hexahistidine tag followed by a tobacco etch virus (TEV) protease cleavage site or a N-terminal StrepII tag followed by a super folder green fluorescent protein (msfGFP) and a tobacco etch virus (TEV) protease cleavage site, respectively.

### Expression and purification of PYROXD1

For expression and purification of PYROXD1, *Escherichia coli* BL21(DE3) cells were transformed with pGEX-6P-1 plasmid encoding for GST-PYROXD1 WT and mutant variants. Bacterial cells were grown at 37°C, 130 rpm in LB medium supplemented with 100 µg ml^−1^ ampicillin. When cultures reached an optical density of 0.6 at 600 nm (OD^600^), temperature was decreased to 18°C and protein expression was induced with 0.1 mM isopropyl-β-D-thiogalactosidase (IPTG), after which cell growth was continued for another 18 hours. Bacterial cells were harvested by centrifugation and flash frozen in liquid nitrogen for storage at -80°C.

Cell pellets were thawed and resuspended in lysis buffer containing 50 mM Tris HCl pH 8.0, 150 mM NaCl, 5 mM MgCl_2_ supplemented with 1 mM AEBSF, 0.1% (v/v) Tween-20, 1 mg ml^−1^ chicken egg white lysozyme, and one proteinase inhibitor tablet per 100 mL (cOmplete Protease Inhibitor, EDTA free). After 30 minutes incubation at 4°C with gentle shaking, cells were lysed by ultrasonication, and the lysate was cleared by centrifugation at 40,000 g for 30 min at 4°C. The cleared lysate was incubated with Glutathione Sepharose beads (Cytiva, GE17-0756-01), equilibrated with lysis buffer, for 2 hours at 4°C with gentle rocking. Next, the beads were washed 5 times with lysis buffer and resuspended in one column volume lysis buffer.

To elute PYROXD1 from the beads, 1 mg of GST-3C Precision protease was added and incubated for 18 hours at 4°C. Subsequently, the beads were loaded onto a gravity flow column and washed with three column volumes of lysis buffer. The collected flowthrough was pooled and concentrated using 30 kDa molecular weight cut-off centrifugal filters (Merck Millipore). The concentrated sample was loaded onto a HiLoad Superdex S75 (16/600) column (Cytiva, 28989333) equilibrated with a buffer containing 20 mM HEPES pH 8.0, 150 mM NaCl and 1 mM TCEP, yielding pure and monodisperse protein. Peak fractions were pooled, concentrated using 30 kDa molecular weight cut-off centrifugal filters (Merck Millipore) and flash-frozen in liquid nitrogen for storage at -80°C.

For the purification of GST-PYROXD1, the protocol described above was altered as follows. After incubating the cleared lysate with Glutathione Sepharose beads, the beads were transferred to a gravity flow column and washed 5 times with lysis buffer. Proteins were eluted from the beads in 5 fractions by the addition one column volume lysis buffer supplemented with 10 mM GSH. Eluted fractions were analyzed by SDS-PAGE and GST-PYROXD1 containing fractions were pooled, after which proteins were dialyzed for 18 hours at 4°C against lysis buffer. The next day the proteins were collected and concentrated using 30 kDa molecular weight cut-off centrifugal filters (Merck Millipore) for size exclusion chromatography.

### Expression and purification of Archease

For expression and purification of Archease, *Escherichia coli* BL21(DE3) Rosetta2 cells were transformed with the 2M-T plasmid encoding for His_6_ -MBP-Archease. Cell cultures were grown at 37°C, 130 rpm in LB medium supplemented with 100 µg ml^−1^ ampicillin. When cultures reached an optical density of 0.6 at 600 nm (OD^600^), the temperature was decreased to 18°C and protein expression was induced with 0.2 mM isopropyl-β-D-thiogalactosidase (IPTG), after which cell growth was continued for another 18 hours. Bacterial cells were harvested by centrifugation and flash frozen in liquid nitrogen for storage at -80°C.

Cell pellets were thawed and resuspended in lysis buffer containing 20 mM Tris pH 8.0, 500 mM NaCl, and 5 mM imidazole, 1 µg ml^−1^ pepstatin and 200 µg ml^−1^ AEBSF. After 30 minutes incubation at 4°C with gentle shaking, cells were lysed by ultrasonication and the lysate was cleared by centrifugation at 40,000g for 30 min at 4°C. The cleared lysate was loaded onto two 5 mL Ni-NTA Superflow cartridges (QIAGEN, 30761) equilibrated with lysis buffer and washed with 10 column volumes of washing buffer containing 20 mM Tris pH 8.0, 500 mM NaCl, and 10 mM imidazole. Next, proteins were eluted with elution buffer containing 20 mM Tris pH 8.0, 500 mM NaCl, and 250 mM imidazole. Protein-containing fractions were pooled and dialyzed for 18 hours at 4°C against 20 mM Tris pH 8.0, 250 mM KCl, 5% glycerol in the presence of His_6_-TEV protease.

To remove the affinity tag, remaining impurities, uncleaved protein and His_6_-TEV protease, the sample was re-applied onto the Ni-NTA Superflow cartridges and the flow-through was pooled and concentrated using 10 kDa molecular weight cut-off centrifugal filters (Merck Millipore). The concentrated sample was loaded onto a HiLoad Superdex S75 (26/600) column (Cytiva, 28989334) equilibrated with a buffer containing 20 mM HEPES pH 8.0, 250 mM KCl and 1 mM DTT, yielding pure and monodisperse protein. Peak fractions were pooled, concentrated using 10 kDa molecular weight cut-off centrifugal filters (Merck Millipore) and flash-frozen in liquid nitrogen for storage at -80°C.

### Expression and purification of RTCB and StrepII-GFP-RTCB

For expression and purification of RTCB and StrepII-GFP-RTCB, recombinant baculoviruses encoding for His_6_-MBP-RTCB or StrepII-GFP-RTCB, respectively, were generated using the Bac-to-Bac Baculovirus expression system (Thermo scientific). These Baculoviruses were used to infect Sf9 insect cells at a density of 1.0·10^6^ ml^-1^. After infection, cells were cultured for 60 hours at 27°C, 90 rpm in SF-4 Baculo Express insect cell culture medium (BioConcept, 9-00F38-N). Insect cells were harvested by centrifugation and flash frozen in liquid nitrogen for storage at -80°C.

To purify RTCB, cell pellets were thawed and resuspended in lysis buffer containing 20 mM Tris pH 8.0, 150 mM NaCl, 5 mM imidazole supplemented with 0.1% Tween 20, and one proteinase inhibitor tablet per 50 mL (cOmplete Protease Inhibitor, EDTA free). Cells were lysed by ultrasonication, and the lysate was cleared by centrifugation at 40,000g for 30 min at 4°C. The cleared lysate was loaded onto Ni-NTA Superflow resin (QIAGEN, 30410) equilibrated with lysis buffer and washed with 10 column volumes of washing buffer containing 20 mM Tris pH 8.0, 500 mM NaCl, and 10 mM imidazole. Next, proteins were eluted with elution buffer containing 20 mM Tris pH 8.0, 500 mM NaCl, and 250 mM imidazole. Protein-containing fractions were pooled and dialyzed for 18 hours at 4°C against 20 mM HEPES pH 8.0, 500 mM KCl, and 3 mM MgCl_2_ in the presence of His_6_-TEV protease.

To remove the affinity tag, remaining impurities, uncleaved protein and His_6_-TEV protease, the sample was re-applied onto Ni-NTA Superflow resin and the flow-through was pooled and concentrated using 10 kDa molecular weight cut-off centrifugal filters (Merck Millipore). The concentrated sample was loaded onto a HiLoad Superdex S200 (16/600) column (Cytiva, GE28-9893-35) equilibrated with a buffer containing 20 mM HEPES pH 8.0, 250 mM KCl and 1 mM DTT, yielding pure and monodisperse protein. Peak fractions were pooled, concentrated using 10 kDa molecular weight cut-off centrifugal filters (Merck Millipore) and flash-frozen in liquid nitrogen for storage at -80°C.

To purify StrepII-GFP-RTCB, cell pellets were thawed and resuspended in lysis buffer containing 20 mM Tris pH 8.0, 150 mM NaCl, supplemented with 0.1% Tween 20, and one proteinase inhibitor tablet per 50 mL (cOmplete Protease Inhibitor, EDTA free). Cells were lysed by ultrasonication and the lysate was cleared by centrifugation at 40,000g for 30 min at 4°C. The clarified lysate was applied to a gravity column containing 5 mL Strep-Tactin resin (IBA, 2-1201-002) and washed with 10 CV wash buffer containing 20 mM HEPES pH 8.0, 500 mM KCl, 1 mM DTT, and 2 mM MgCl_2_. Proteins were eluted with wash buffer supplemented with 2.5 mM desthiobiotin. Eluted proteins were pooled and concentrated using 10 kDa molecular weight cut-off centrifugal filters (Merck Millipore). The concentrated sample was loaded onto a HiLoad Superdex S200 (16/600) column (Cytiva, GE28-9893-35) equilibrated with a buffer containing 20 mM HEPES pH 8.0, 500 mM KCl, 2 mM MgCl_2_, and 1 mM DTT, yielding pure and monodisperse protein. Peak fractions were pooled, concentrated using 10 kDa molecular weight cut-off centrifugal filters (Merck Millipore) and flash-frozen in liquid nitrogen for storage at -80°C.

### RTCB-PYROXD1 complex sample preparation and cryo-EM data collection

Prior to grid preparation for cryo-EM, thawed protein samples of RTCB and PYROXD1 were incubated at a 1:1 molar ratio by mixing 100 µM RTCB with 100 µM PYROXD1 in 100 µL buffer containing 20 mM HEPES pH 8.0, 150 mM NaCl, 0.5 mM TCEP, 5 mM MgCl_2_ and 0.5 mM NADH. After 30 minutes of incubation on ice, samples were loaded onto a Superdex 200 (10/300) size-exclusion chromatography column (GE Healthcare) equilibrated with a buffer containing 20 mM HEPES pH 8.0, 150 mM NaCl, 0.5 mM TCEP, 5 mM MgCl_2_ and 0.5 mM NADH (Sigma-Aldrich, N8129). Protein complex-containing fractions were collected and concentrated using 10 kDa molecular weight cut-off centrifugal filters (Merck Millipore). The concentrated protein was split into two samples, a sample at a protein concentration of 1.5 mg mL^-1^ that was supplemented with 0.01% Octyl-beta-Glucoside (dataset 1) and a sample at a protein concentration of 0.6 mg mL^-1^ (dataset 2). To each 200-mesh holey carbon grid (Au R1.2/1.3, Quantifoil Micro Tools), 2.5 µl of sample was applied and blotted for 4 s at 80% humidity and 4 °C. Grids were plunge frozen in liquid ethane using a Vitrobot Mark IV plunger (Thermo Scientific), and stored in liquid nitrogen until cryo-EM data collection. Cryo-EM data collection was performed on a FEI Titan Krios microscope equipped with a Gatan K3 direct electron detector (University of Zurich) operated at 300 kV in super-resolution counting mode. Data acquisition was performed using the EPU Automated Data Acquisition Software for Single Particle Analysis from ThermoFisher with three shots per hole at defocus range of −1.0 μm to −2.4 μm (0.2-µm steps). The final datasets comprised a total of 14,516 micrographs (dataset 1: 5,114 micrographs & dataset 2: 9,302 micrographs) at a calibrated magnification of 130,000x and a in super-resolution pixel size of 0.325 Å. Micrographs for dataset 1 were exposed for 1.01 s with a total dose of 64.592 e^−^ Å^−2^ over 38 subframes, whereas the micrographs for dataset 2 were exposed for 1.01s with a total dose of 66.459 e^−^ Å^−2^ over 38 subframes.

### Cryo-EM data processing and model building

Cryo-EM data was processed using cryoSPARC (v3.3.2)^31^. The total of 14,516 micrographs were imported and motion-corrected with patch motion correction (multi) after which the CTF values of the micrographs were estimated using patch CTF estimation (multi). Micrographs were curated and micrographs with a resolution estimate >5 Å and a defocus >2.4 µm were discarded from the dataset (742 micrographs), yielding 13,774 micrographs for further processing steps. Next, an initial set of particles was picked with blob picker using an elliptical blob and a minimum and maximum particle diameter of 50 and 150 Å, respectively. After extraction of the particles with a box size of 288×288 pixels, particles were subjected to 2D classification to generate templates for picking (5 templates). After template-based picking with a particle diameter of 100 Å, particles were extracted and subjected to 2D classification with a circular mask of 110 Å. Classes with defined particles were selected, resulting in a total of 3,097,252 particles, which were used to generate one ab initio model which was used for heterogeneous refinement with four classes. Classes were inspected visually using UCSF Chimera^32^, and the particles and volume of the best class were subjected to homogenous refinement. The output was used for variability analysis with 3 modes, filtered at a resolution of 5 Å. After visual inspection of the variability analysis using UCSF Chimera^32^, two volumes were used for heterogeneous refinement with four classes. Classes were inspected, and the particles and volume of the best class were subjected to non-uniform refinement with optimization of CTF parameters enabled. The final map was sharpened with a B-factor of -140. The local resolution was estimated based on the resulting map using the local resolution function of cryoSPARC and displayed on the map using UCSF Chimera^32^. The structural model of the RTCB-PYROXD1 was built in Coot (V0.9.2)^33^ and was refined over multiple rounds using Phenix^34,35^. Real-space refinement was performed with the global minimization, atomic displacement parameter (ADP) refinement and secondary structure restrains enabled. The quality of the atomic model, including protein geometry, Ramachandran plots, clash analysis and model cross-validation, was assessed with MolProbity and the validation tools in Phenix^34–37^. The refinement statistics of the final model are listed in Table S1. Figures of maps, models and the calculations of map contour levels were generated using UCSF Chimera^32^.

### In vitro pull-down assays of RTCB-PYROXD1 complexes

To immobilize RTCB, 0.2 nmol of StrepII-GFP-RTCB was diluted in 200 μL of binding buffer containing 20 mM HEPES pH 8.0, 150 mM KCl, 0.1% Tween-20, 1 mM TCEP, 2.5 mM MgCl_2_, 2.5 mM MnCl_2_ and 0.5 mM NADH (Sigma-Aldrich, N8129). This solution was incubated by gently rocking for 30 min at 4°C with 2 μL (packed volume) of MagStrep “type 3” XT beads (IBA, 2-4090-002). The supernatant was removed and the beads were washed twice with 500 μL of binding buffer. Next, 0.4 nmol of PYROXD1 in 200 μL of reaction buffer containing HEPES pH 8.0, 150 mM KCl, 0.1% Tween-20, 1 mM TCEP, 0.5 mM NADH (Sigma-Aldrich, N8129) supplemented with 2.5 mM MgCl_2_ and 2.5 mM MnCl_2_ was added to the washed beads and incubated gently by rocking for 30 min at 4°C. To test the influence of metals on PYROXD1 binding, the reaction buffer was supplemented with either 5 mM (MgCl_2_, MnCl_2_, NiCl_2_, CoCl_2_, CaCl2) or 0.1 mM (ZnCl_2_, CuCl_2_) of divalent metal. After incubation, the beads were washed three times with 500 μL of reaction buffer. After washing, the beads were resuspended in 40 μL of 1x SDS loading dye (45 mM Tris pH 6.8, 10% glycerol, 1% SDS, 50 mM DTT, 0.002% bromophenol blue). The samples were incubated at 95°C for 5 min and 10 μL of the sample was resolved on a 4%–15% Mini-PROTEAN® TGX Precast Gel (Biorad) and stained with Coomassie Blue.

### RNA ligation assays with PYROXD1 variants

Ligation assays were carried out following a two-step procedure. In the first step, the oxidation step, 29 nM of recombinant tRNA-LC (containing RTCB, DDX1, CGI-99 and FAM98b) was incubated with 290 nM Archease, 100 nM PYROXD1 and H_2_O_2_ at various concentrations for 1 h at 20°C at 600 rpm in a reaction buffer containing 30 mM HEPES/KOH (pH 7.4), 100 mM KCl, 5 mM MgCl_2_, 0.1 mM AEBSF, 10% glycerol, 1% NP40, which was supplemented with 440 µM NADPH. Separately, a solution containing 12.5 mM TCEP, 100 mM KCl, 2.9 mM MgCl_2_, 7.5 mM ATP, 0.5 mM GTP, 250 mM ZnCl_2_, RNasin®Ribonuclease Inhibitors (1 mL per 500 mL) and 65% (v/v) glycerol was mixed (2/1, v/v) with the radiolabelled oligoribonucleotide to generate a reaction cocktail. The reaction cocktail was then mixed with the samples from the oxidation step in a 3 to 2 (v/v) ratio, respectively. In the second step of the procedure, the ligation step, samples were incubated for 45 min at 30°C. The final concentrations of the critical components in the ligation step were: 12 nM tRNA-LC, 120 nM Archease, 3.2 mM ATP, 230 µM GTP, 180 µM NADPH and 40 nM PYROXD1. The ligation reactions were then quenched with 2% Formamide solution (1/1, v/v), composed of 90% Formamide, 50 mM EDTA, 1 ng/ mL bromophenol blue, 1 ng/ml Xylene Cyanol and incubated for 1 min at 95°C. The reactions were resolved on a 15% denaturing polyacrylamide gel and detected by autoradiography using a Typhoon 5 phosphorimager.

The levels of oxidative inactivation of the tRNA-LC are dependent on the ratio of NAD(P) H/NAD(P)+, presence trace metal ions and the concentration of reducing agents in the reaction buffer. To obtain reproducible results it is important that NAD(P)H stocks are prepared freshly prior to the experiment from NAD(P)H that has not been stored long-term. Additionally, the assay is sensitive to the levels of trace metals that are present in the water, hence buffers need to be prepared in bulk, aliquoted and stored at -20°C until use.As tRNA-LC is prepared in a buffer with reducing agent, it is important to maintain a constant dilution factor between experiments to account for the presence of residual reducing agent in the solution. Lastly, fresh glycerol should be used for the assays and storage, as glycerol that has been exposed light and high temperatures may contain traces of electrophilic dehydration/oxidation products that inactivate tRNA-LC.

### In vitro competition assays

For competition assays, 0.2 nmol of StrepII-GFP-RTCB was diluted in 200 μL of binding buffer. This solution was incubated by gently rocking for 30 min at 4°C with 2 μL (packed volume) of MagStrep “type 3” XT beads (IBA, 2-4090-002). The supernatant was removed and the beads were washed twice with 500 μL of binding buffer. Next, 0.4 nmol of PYROXD1 and 2 nmol Archease in 200 μL of reaction buffer containing HEPES pH 8.0, 150 mM KCl, 0.1% Tween-20, 1 mM TCEP, 2.5 mM MgCl_2_, 2.5 mM MnCl_2_ supplemented with 0.5 mM NADH (Sigma-Aldrich, N8129), 1 mM GTP (Roth, K056.4) or 1 mM GMP (Jena Biosciences, JBS-NU-1028), was added to the washed beads and incubated gently by rocking for 30 min at 4°C. After incubation, the beads were washed three times with 500 μL of reaction buffer. After washing, the beads were resuspended in 40 μL of 1x SDS loading dye (45 mM Tris pH 6.8, 10% glycerol, 1% SDS, 50 mM DTT, 0.002% bromophenol blue). The samples were incubated at 95°C for 5 min and 10 μL of the sample was resolved on a 4%–15% Mini-PROTEAN® TGX Precast Gel (Biorad) and stained with Coomassie blue dye.

### Spectroscopic assays

For spectroscopic analysis of NADH oxidation in pre-steady state conditions, PYROXD1 was diluted to 25 µM in 100 μL of buffer containing 20 mM Tris-HCl pH 8.0, 150 mM KCl, 0.05% Tween-20. Diluted samples were transferred to a 96-well plate (Corning, 3635) and placed into a PHERAstar FSX plate reader (BMG Labtech). Next, NADH (Sigma-Aldrich, N8129) was injected into the wells to a final concentration of 25 µM and absorbance at 340 nm was measured over 30 minutes with 3-minute intervals at room temperature. For control experiments, the absorbance at 340 nm of 25 µM NADH (Sigma-Aldrich, N8129), 25 µM FAD (Sigma-Aldrich, F6625) and a mixture of 25 µM NADH and 25 µM FAD were tracked over 30 minutes with 3-minute intervals at room temperature. For spectroscopic analysis of NADH oxidation under steady-state conditions, PYROXD1 was diluted to 2.5 µM in 100 μL of buffer containing 20 mM Tris-HCl pH 8.0, 150 mM KCl, 0.05% Tween-20. Diluted samples were transferred to a 96-well plate (Corning, 3635) and placed into a PHERAstar FSX plate reader (BMG Labtech). Next, NADH (Sigma-Aldrich, N8129) was injected into the wells to a final concentration of 250 µM and absorbance at 340 nm was measured over 30 minutes with 3-minute intervals at room temperature.

### Co-precipitation of guanylylated RTCB

For the guanylylation reaction, 4 nmol of RTCB was incubated with 20 nmol Archease in 200 μl guanylylation buffer containing 20 mM HEPES pH 8.0, 100 mM NaCl, 5 mM DTT, 2.5 mM MnCl_2_, 0.5 mM GTP (Roth, K056.4) for 60 min at 25°C. After the incubation, 25 μl of the sample were removed, flash frozen in liquid nitrogen and stored at -80°C until subsequent analysis by liquid-chromatography-mass spectrometry (LC-MS). In parallel, 25 μl (packed volume) of Glutathione Sepharose™ 4 Fast Flow beads (Sigma, GE17-5132-01) were equilibrated in guanylylation buffer, after which the beads were incubated with 1 nmol GST-PYROXD1 in 200 μl of guanylylation buffer for 30 min at 4°C, gently rocking. Subsequently, the beads were washed two times with 200 μl guanylylation buffer and added to the guanylylation sample. After supplementing the mixture with 0.5 mM NADH (Sigma-Aldrich, N8129), the sample was incubated for 15 min at 4°C, gently rocking. After the incubation, the sample was centrifuged for 2 min at 500x g and 4°C, and the supernatant was recovered, flash frozen in liquid nitrogen and stored at -80°C for subsequent analysis by LC-MS. Next, the beads were washed three times with 500 μl guanylylation buffer supplemented with 0.5 mM NADH, followed by elution of the proteins with 40 μl of elution buffer containing 20 mM HEPES pH 8.0, 150 mM KCl 10 mM GSH, 2.5 mM MnCl_2_. The eluted proteins were flash frozen in liquid nitrogen and stored at -80°C for subsequent analysis by LC-MS. The experiment was carried out in triplicate.

### Quantification of guanylylated RTCB by LC-MS

To quantify the levels of guanylylated RTCB by LC-MS, samples were diluted 1:3 with water after wich 5 μl was injected into an ACQUITY UPLC™ BioResolve-RP-mAb polyphenyl column (450 angstrom, 2.7 μm, 2.1 mm x150 mm) column (Waters, USA). To separate proteins, a gradient was applied from 0.1 % Difluoroacetic acid (DFA) in water to 0.1 % DFA in Acetonitril/75 % 2-propanol at 60 min over 15 min. LC-MS analysis was performed on a SYNAPT G2-Si mass spectrometer coupled to an ACQUITY UPLC station.

### Ligation assays to assess the effects of guanylation on oxidative inactivation of RTCB

These experiments were carried out using a three-step procedure. In the first step, the guanylylation step, 98 nM of recombinant tRNA-LC (containing RTCB, DDX1, CGI-99 and FAM98b) was incubated for 1 h at 30°C, 600 rpm with 980 µM ATP and where designated 980 nM Archease and/or 220 µM GTP in a reaction buffer containing 30 mM HEPES/KOH (pH 7.4), 100 mM KCl, 5 mM MgCl_2_, 0.1 mM AEBSF, 10% glycerol, and 1% NP40. Next, in the second step, the oxidation step, NADPH, H_2_O_2_ and PYROXD1 were added, resulting in the following concentrations of the critical components: 29 nM tRNA-LC, 290 nM archease, 290 µM ATP, 67 µM GTP, 440 µM NADPH, 100 nM PYROXD1 and the designated concentrations of H_2_O_2_. Subsequently, solutions of GTP or Archease were added to supplement the for component missing in the guanylylation and oxidation steps to ensure that all reactions enter the final ligation step with the same reagent compositions. Ligation step was carried out and the products were analysed as described above.

### Dissociation kinetics analysis of RTCB-PYROXD1

For the RTCB dissociation assay, 2 nmol 2xStrepII-GFP-RTCB and 4 nmol PYROXD1 were diluted in 3 mL binding buffer containing 150 mM KCl, 20 mM HEPES-KOH pH 8, 1 mM TCEP, 0.1% Tween 20, 5 mM MgCl_2_, 0.5 mM NADH and incubated for 10 min on ice. Next, 400 µL slurry of Magnetic Streptactin beads (MagStrep “type3” XT beads – IBA 2-4090-002) equilibrated with binding buffer and added to the 2xStrepII-GFP-RTCB and PYROXD1 mixture followed by incubation at 1 h at 4.C, rocking gently. Subsequently, the supernatant was removed and the magnetic beads were transferred to a 1.5 ml reaction tube. After washing the beads twice with 1 mL wash buffer containing 150 mM KCl, 20 mM HEPES-KOH pH 8, 1 mM TCEP, 0.1% Tween 20, 5 mM MgCl_2_, the resin was equally distributed to multiple 1.5 mL reaction and tubes and the resin resuspended in 400 µL wash buffer. At the indicated time points the supernatant was removed, the resin washed twice with 400 µL wash buffer and the beads were resuspended in 40 µL SDS sample buffer.

