## Supplementary Figures S1-4, Supplementary Table S1 for "Mechanistic basis for oxidative stress protection of the human tRNA ligase complex by the oxidoreductase PYROXD1"

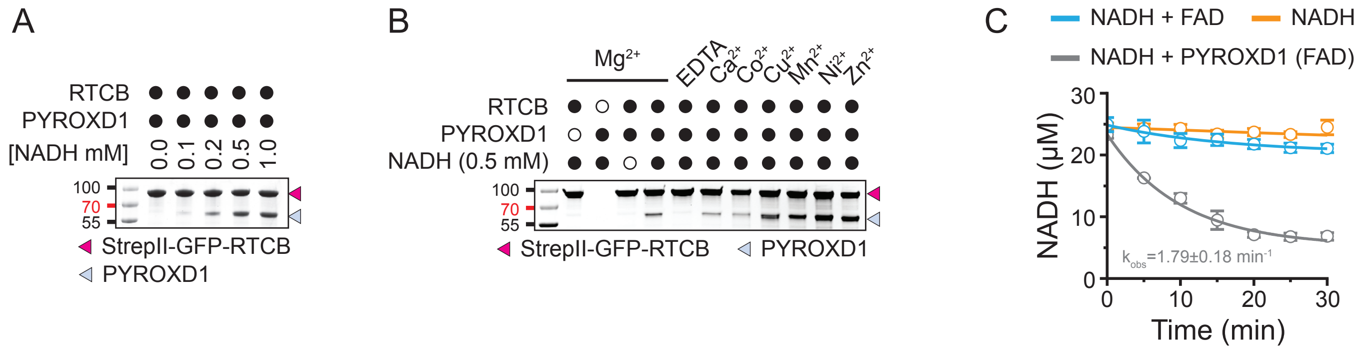


**Figure S1: RTCB-PYROXD1 complex formation and spectroscopic analysis of NADH turnover by PYROXD1, related to Figure 1.**

**(A)** In vitro pull-down experiment with recombinant PYROXD1 and immobilized StrepII-GFP-RTCB in the presence of varying concentrations of NADH. Strep-Tactin beads were washed to remove unbound PYROXD1, and bound proteins were analyzed by SDS-PAGE and Coomassie blue staining. **(B)** In vitro pull-down experiment with PYROXD1 mutants and StrepII-GFP-RTCB in the presence of divalent metal ions. Strep-Tactin beads were washed to remove unbound PYROXD1, and bound proteins were analyzed by SDS-PAGE and Coomassie blue staining. **(C)** Spectroscopic analysis of NADH oxidation by human PYROXD1. Data points represent the mean ± SEM of three independent replicates. Solid lines represent a single-exponential fit.

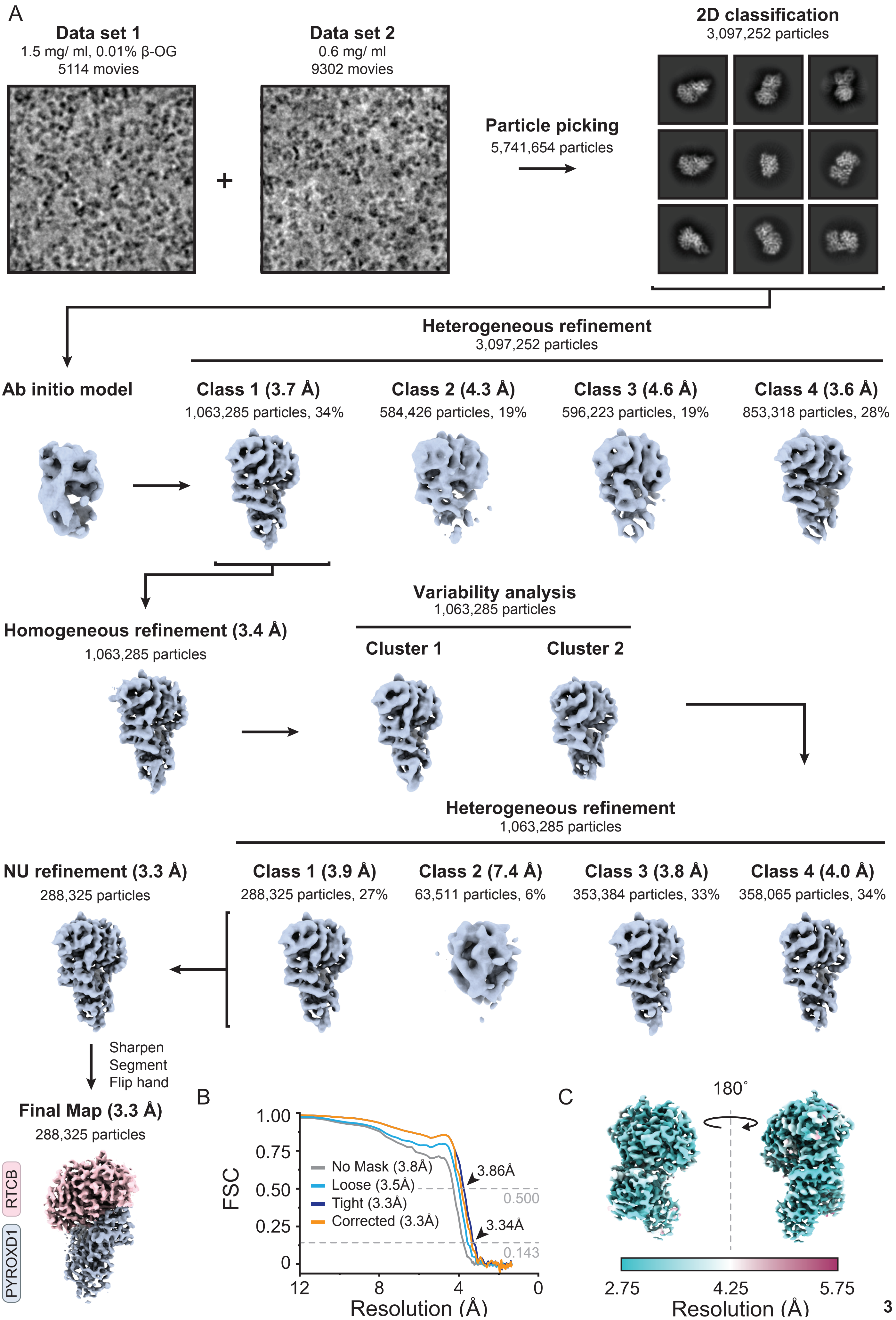
A
Data set 1
1.5 mg/ ml, 0.01% β-OG
5114 movies
Data set 2
0.6 mg/ ml
9302 movies
Particle picking
5,741,654 particles
2D classification
3,097,252 particles
Heterogeneous refinement
3,097,252 particles
Ab initio model
Class 1 (3.7 Å)
1,063,285 particles, 34%
Class 2 (4.3 Å)
584,426 particles, 19%
Class 3 (4.6 Å)
596,223 particles, 19%
Class 4 (3.6 Å)
853,318 particles, 28%
Homogeneous refinement (3.4 Å)
1,063,285 particles
Variability analysis
1,063,285 particles
Cluster 1
Cluster 2
Heterogeneous refinement
1,063,285 particles
NU refinement (3.3 Å)
288,325 particles
Class 1 (3.9 Å)
288,325 particles, 27%
Class 2 (7.4 Å)
63,511 particles, 6%
Class 3 (3.8 Å)
353,384 particles, 33%
Class 4 (4.0 Å)
358,065 particles, 34%
Sharpen
Segment
Flip hand
Final Map (3.3 Å)
288,325 particles
RTCB
PYROXD1
B
FSC
No Mask (3.8Å)
Loose (3.5Å)
Tight (3.3Å)
Corrected (3.3Å)
3.86Å
3.34Å
0.500
0.143
Resolution (Å)
C
180˚
2.75
4.25
5.75
Resolution (Å)

**Figure S2: Cryo-EM processing workflow for the human RTCB-PYROXD1 complex, related to Figure 1.**

**(A)** Cryo-EM processing workflow for the human RTCB-PYROXD1 complex. **(B)** Fourier Shell Correlation (FCS) determined from two independently refined half maps. The gold standard cut-off (FCS=0.143) is marked with an arrow). **(C)** Local resolution estimation on the final cryo-EM density map of the human RTCB-PYROXD1 complex.

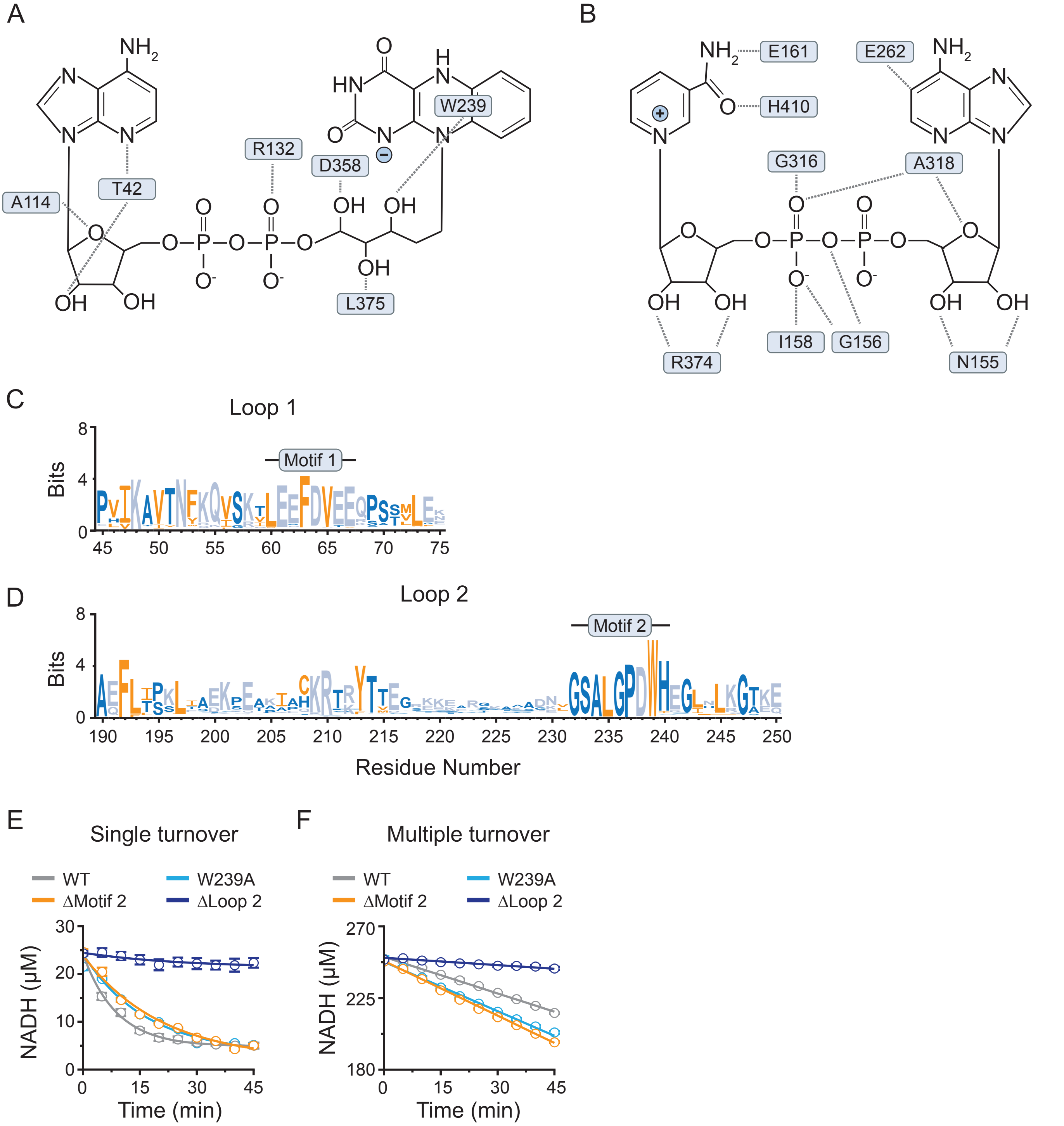


**Figure S3: Loops in PYROXD1 allosterically control its activity, related to Figure 1 and 3.**

**(A)** Interaction map of the FADH⁻ ligand in the RTCB-PYROXD1 complex. **(B)** Interaction map of the NAD⁺ ligand in the RTCB-PYROXD1 complex. **(C)** Sequence conservation analysis of loop 1 in PYROXD1. Figure was generated using WebLogo[28]. Black line indicates a conserved sequence motif (motif 1) in loop 1. **(D)** Sequence conservation analysis of loop 2 in PYROXD1. Black line indicates a conserved sequence motif (motif 2) in loop 2. **(E)** Single-turnover kinetics of NADH oxidation by human PYROXD1 variants under aerobic conditions. Data points represent the mean ± SEM of three independent replicates. Solid lines represent a single-exponential fit. **(F)** Multiple-turnover kinetics of NADH oxidation by human PYROXD1 variants under aerobic conditions. Data points represent the mean ± SEM of three independent replicates. Solid lines represent a linear fit.

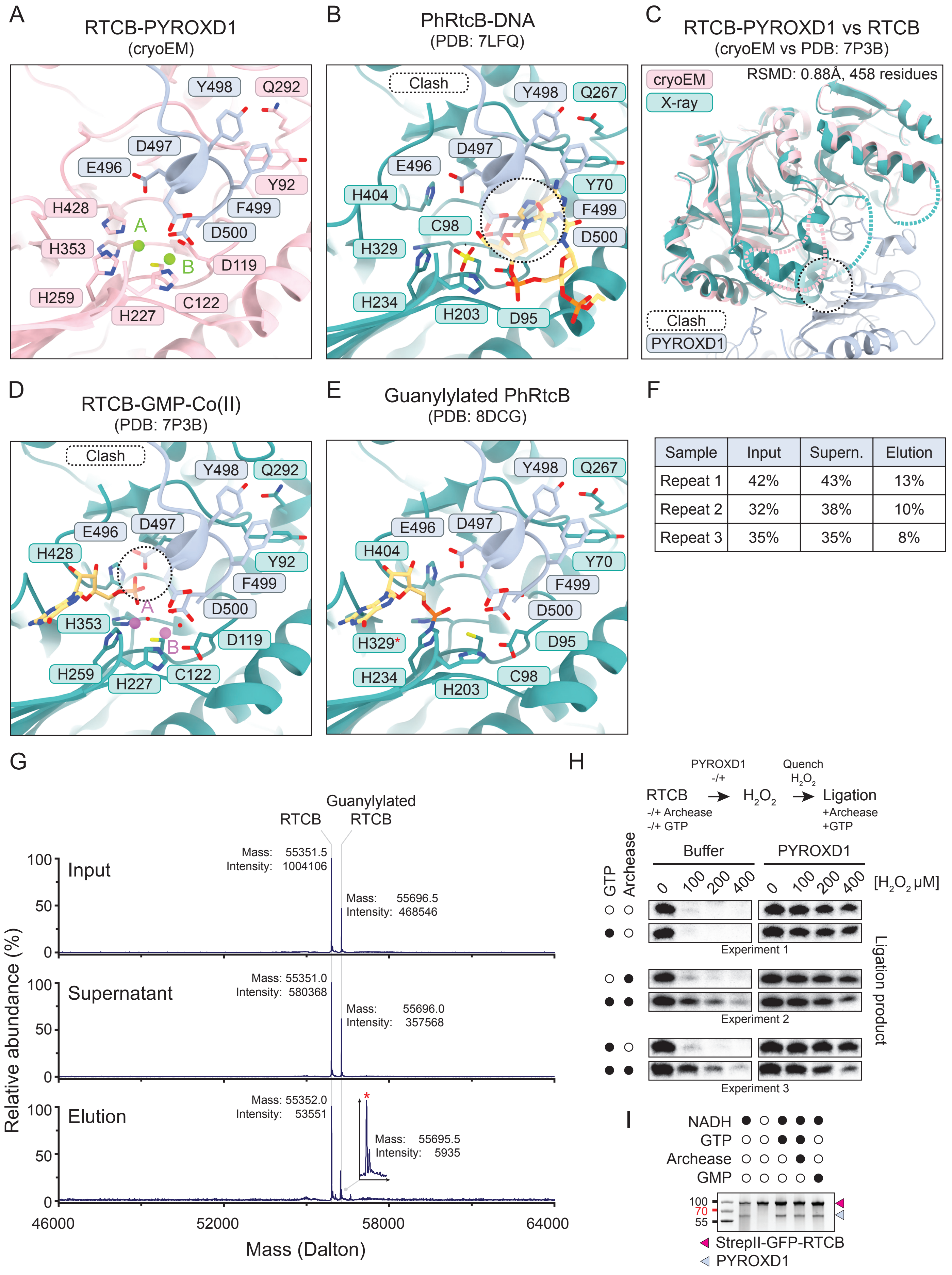


| Sample | Input | Supern. | Elution |
|---|---|---|---|
| Repeat 1 | 42% | 43% | 13% |
| Repeat 2 | 32% | 38% | 10% |
| Repeat 3 | 35% | 35% | 8% |

**Figure S4: Structural comparison of RTCB catalytic centers, related to Figure 4.**

**(A)** Detailed view of the interaction of the CTD of PYROXD1 with the catalytic center of RTCB. Bound magnesium ions are depicted as green spheres. **(B)** Structural superposition of the PYROXD1 CTD onto catalytic cleft of

DNA-bound PhRtcB (PDB: 7LFQ)[25]. Steric clashes are indicated by a dotted circle. **(C)** Structural superposition of the PYRXOD1-RTCB complex and GMP-bound RTCB (PDB: 7P3B)[9]. **(D)** Structural superposition of the PYROXD1 CTD with the catalytic center of GMP-bound human RTCB (PDB: 7P3B)[9]. Clashes are indicated by a dotted circle. **(E)** Structural comparison of the position of the CTD of PYROXD1 in the catalytic center of guanlylylated PhRtcB (PDB: 8DCG)[29]. **(F)** Fraction of guanlylated RTCB in pull-down samples, as determined by LC-MS. Guanylylation levels are indicated for three independent replicates. **(H)** Representative mass spectra of the input, unbound (supernatant) and bound (elution) fractions. Mass and intensity of each peak are indicated. Red asterisk indicates a mass peak corresponding to GSH-modified RTCB. **(I)** In vitro ligation assay to determine the effects of guanylylation on the oxidative inactivation of RTCB. The ligation assay was performed in a three-step procedure, where RTCB was incubated with GTP and Archease (see labels on the left), followed by incubation with varying concentrations of $H_2O_2$ in the presence or absence of PYROXD1. In the last steps all samples were supplemented with GTP and Archease to allow for multiple-turnover ligation of an RNA substrate.

**Table S1: Cryo-EM data collection, refinement and validation statistics**

| | Human RTCB-PYROXD1 complex (EMDB-xxxx) (PDB xxxx) |
|---|---|
| **Data collection and processing** | |
| Magnification | 130,000 |
| Voltage (kV) | 300 |
| Electron exposure (e–/Å$^2$) | Data set 1: 64.592<br>Data set 2: 63.163 |
| Defocus range (μm) | -1.0 to -2.4 (-0.2 steps) |
| Pixel size (Å) | 0.65 |
| Symmetry imposed | C1 |
| Initial particle images (no.) | 3,097,252 |
| Final particle images (no.) | 288,325 |
| Map resolution (Å) | 3.3 |
| FSC threshold | 0.143 |
| Map resolution range (Å) | 2.75-5.75 |
| **Refinement** | |
| Initial model used (PDB code) | 7P3B & 6ZK7 |
| Model resolution (Å) | 3.3 |
| FSC threshold | 0.143 |
| Model resolution range (Å) | 3.3-3.9 |
| Map sharpening *B* factor (Å$^2$) | -140 |
| Model composition | |
| Non-hydrogen atoms | 7291 |
| Protein residues | 891 |
| Ligands | FDA 1<br>NAD 1<br>MG 2 |
| *B* factors (Å$^2$) | |
| Protein | 123.04/373.70/197.80 |
| Ligand | 142.44/192.50/185.46 |
| R.m.s. deviations | |
| Bond lengths (Å) | 0.013 |
| Bond angles (°) | 1.188 |
| Validation | |
| MolProbity score | 2.51 |
| Clashscore | 67.11 |
| Poor rotamers (%) | 0.41 |
| Ramachandran plot | |
| Favored (%) | 96.71 |
| Allowed (%) | 3.29 |
| Disallowed (%) | 0.00 |
